# Hemangiosarcoma Cells Promote Conserved Host-Derived Hematopoietic Expansion

**DOI:** 10.1101/2021.05.21.445198

**Authors:** Jong Hyuk Kim, Ashley J. Schulte, Aaron L. Sarver, Mathew G. Angelos, Aric M. Frantz, Colleen L. Forster, Timothy D. O’Brien, Ingrid Cornax, M. Gerard O’Sullivan, Nuojin Cheng, Mitzi Lewellen, LeAnn Oseth, Sunil Kumar, Susan Bullman, Chandra Sekhar Pedamallu, Sagar M. Goyal, Matthew Meyerson, Troy C. Lund, Jessica Alfoldi, Matthew Breen, Kerstin Lindblad-Toh, Erin B. Dickerson, Dan S. Kaufman, Jaime F. Modiano

## Abstract

Hemangiosarcoma and angiosarcoma are soft-tissue sarcomas of blood vessel-forming cells in dogs and humans, respectively. These vasoformative sarcomas are aggressive and highly metastatic, with disorganized, irregular blood-filled vascular spaces. Our objective was to define molecular programs which support the niche that enables progression of canine hemangiosarcoma and human angiosarcoma. Dog-in-mouse hemangiosarcoma xenografts recapitulated the vasoformative and highly angiogenic morphology and molecular characteristics of primary tumors. Blood vessels in the tumors were complex and disorganized, and they were lined by both donor and host cells, a trait that was not observed in xenografts from canine osteosarcoma and lymphoma. In some cases, the xenografted hemangiosarcoma cells created exuberant myeloid hyperplasia and gave rise to lymphoproliferative tumors of mouse origin. We did not uncover a definitive transmissible etiology, but our functional analyses indicate that hemangiosarcoma cells generate a microenvironment that supports expansion and differentiation of hematopoietic progenitor populations. We conclude that canine hemangiosarcomas, and possibly human angiosarcomas, originate from stromal cells that are part of the bone marrow niche and that these cells may also support the growth of hematopoietic tumors.

**Significance:** We demonstrate that molecular programs supporting expansion of immune and inflammatory cells in hemangiosarcoma resemble those of bone marrow niche cells, providing insights into the potential roles of these cells - whether physiological or pathological - in creating a permissive environment for the progression of hematopoietic malignancies.

## Introduction

Canine hemangiosarcoma, a vasoformative tumor originating from bone marrow (BM)-derived progenitor cells, is a common occurrence in dogs, unlike the rare human angiosarcoma (1–4). Tumors from both species exhibit similar histology and natural history of disorganized, tortuous, and dilated blood vessels with high proliferative activity and metastatic potential (5,6). Molecularly, convergent transcriptional programs involving angiogenesis and inflammation are observed in both canine hemangiosarcoma and human angiosarcoma (7,8). While the tumor microenvironment plays a crucial role in tumor cell survival, disease progression, and metastasis through the niche it creates (9), its contribution to the molecular programs of hemangiosarcoma remains incompletely understood. Recent evidence suggests bidirectional interactions between cancer cells and niche cells, where cancer cells re-educate niche cells and vice versa, particularly in maintaining stemness and self-renewal of cancer stem cells in various tumors (9–11), including hematopoietic tumors (12–14).

The hematopoietic niche is composed of osteoblasts and various stromal cells including endosteal, endothelial cells, fibroblasts, nestin-positive mesenchymal stromal cells, leptin receptor-positive stromal cells, and CXCL12-abundant reticular (CAR) cells. These cells support hematopoiesis by regulating the proliferation and differentiation of hematopoietic stem and progenitor cells (HSPCs) into lineage-committed cells (15). Dysregulation of HSPC function can result in blood disorders (16,17) and hematopoietic malignancies such as leukemia and lymphoma (18–20).

In the present study, we aimed to elucidate the cellular and molecular origin of canine hemangiosarcoma. Our findings indicate that canine hemangiosarcoma cells not only have the capability to form vasoformative tumors but also create a niche for expansion and differentiation of blood cells, supporting an ontogenetic derivation from bone marrow stromal cells.

## Results

### Canine hemangiosarcoma cells can recapitulate the disease *in vivo*

We used *in vivo* xenografts to investigate the biological behavior of canine hemangiosarcoma (21–25). Mice were inoculated with canine hemangiosarcoma cell lines or primary tissues, as detailed in **Supplementary Table S1**. However, the engraftment efficiency of canine hemangiosarcoma xenografts was found to be low. Only cultured DHSA-1426 cells, when injected subcutaneously, consistently generated vasoformative tumors in immunodeficient beige-nude-xid (BNX) mice (**Figure 1**). The tumorigenic potential of this cell line was maintained over multiple passages, as demonstrated by serial passaging of cells cultured from tumor xenografts into new recipient mice (**Supplementary Figure S1**).

**Figure 1.**
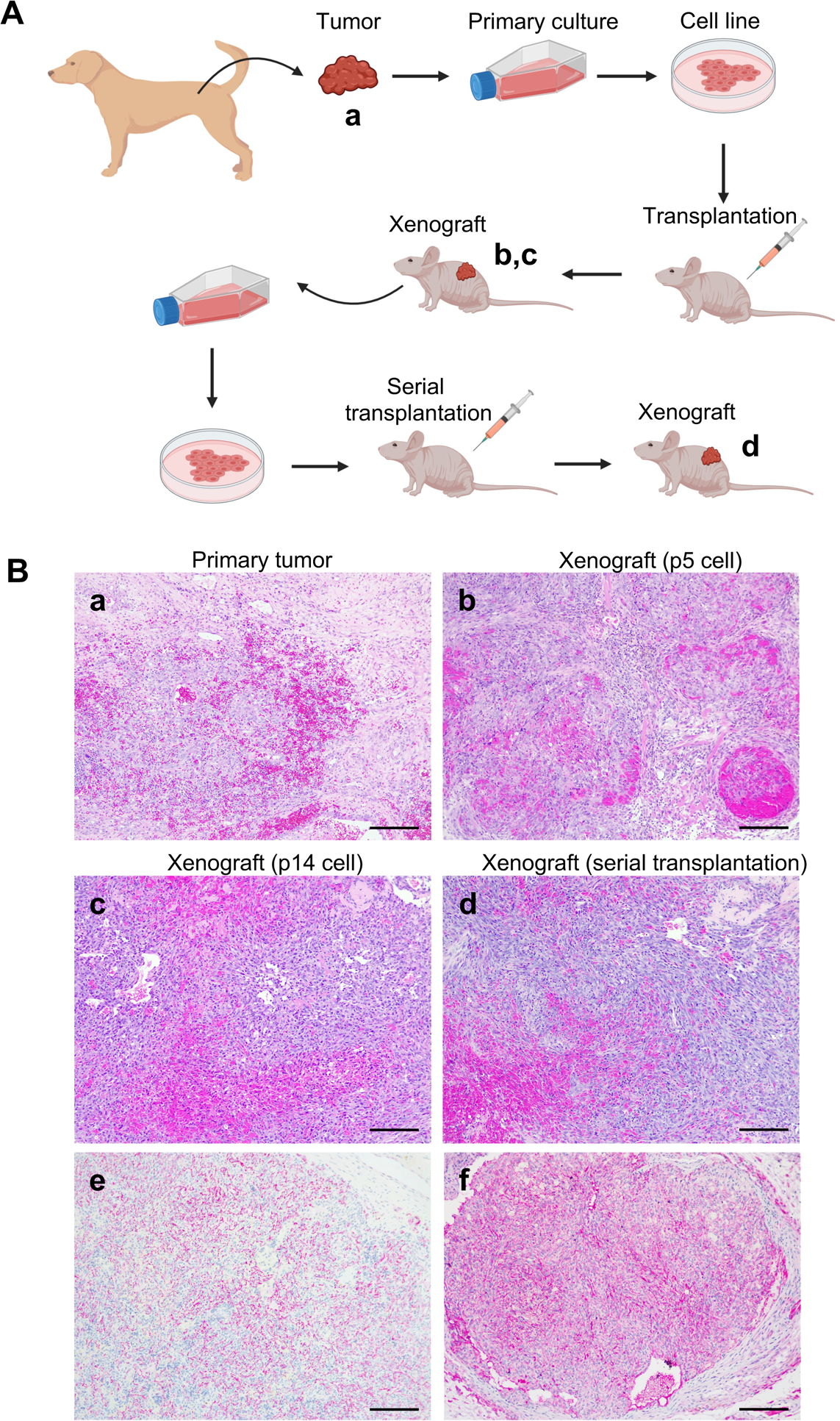
Establishment of xenografts derived from canine hemangiosarcoma in immunodeficient mice. (**A**) Schematic illustration depicts process of tumor xenografts in beige-nude-xid (BNX) mice. (**B**) DHSA-1426 tumor cell line was established from a canine patient diagnosed with hemangiosarcoma (**a**). Cells from DHSA-1426 passages 5 (p5) and 14 (p14) formed tumors histologically classified as hemangiosarcoma (**b, c**). Cells cultured from xenograft-derived tumors developed histologically identical tumors after serial transplantation (**d**). Both the primary tumor (**e**) and the xenograft tumor (**f**) were positive for CD31 immunohistochemical staining. (a-d) Hematoxylin and eosin (H&E) stain. (e,f) Immunohistochemistry with an anti-CD31 antibody (alkaline phosphatase conjugates; counterstain = hematoxylin). Bar = 200 µm.

### Xenografts allow quantification of the stromal contribution in the hemangiosarcoma microenvironment

To establish the contribution of stromal elements to the formation of canine hemangiosarcoma, we utilized fluorescence *in situ* hybridization (FISH) to quantify the presence of both canine and mouse cells in the tumors. Specifically, we employed probes for canine *CXCL8*, as this gene is absent in the mouse genome, and a unique region of the mouse X chromosome, to differentiate between canine and mouse cells, respectively. Our data revealed that the hemangiosarcoma xenografts displayed a complex topological organization, with blood vessels lined by both donor and host cells (**Figure 2A**). Notably, the organization pattern observed in canine hemangiosarcoma xenografts differed from that of canine osteosarcoma and lymphoma xenografts (**Figure 2B**). The hemangiosarcoma tumors were comprised of a mixture of 50-70% malignant canine cells and 30-50% mouse stromal cells. The cellular composition of orthotopic osteosarcoma xenografts was comparable, but without malignant canine cells lining blood vessels. In contrast, lymphoma xenografts exhibited a markedly distinct composition, with less than 5% mouse stromal cells and the absence of malignant canine cells from blood vessels **(Figure 2C**). These findings highlight the utility of xenografts as a valuable tool for quantifying the contribution of stromal elements to the tumor microenvironment.

**Figure 2.**
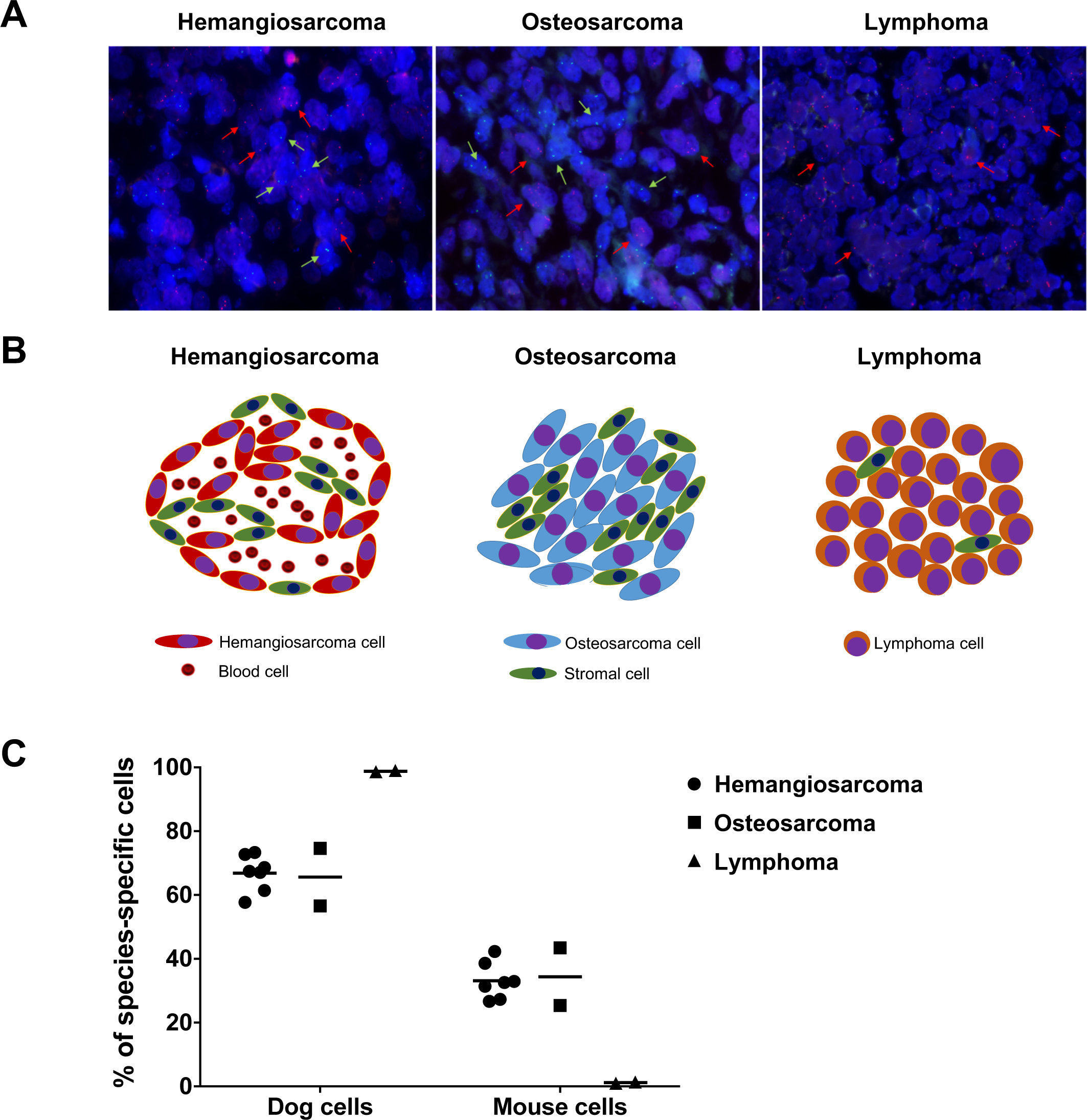
Organization of tumor and stromal cells in mouse xenografts of canine hemangiosarcoma, osteosarcoma, and lymphoma. (**A**) Fluorescence *in situ* hybridization images using canine-specific (*CXCL8*, red) and mouse-specific (X chromosome, green) probes in a canine hemangiosarcoma xenograft, a canine osteosarcoma xenograft, and a canine lymphoma xenograft transplanted into receptive immunodeficient female mouse hosts. Red and green arrows point to representative xenograft canine tumor cells and mouse stromal cells, respectively, to aid in identification. (**B**) Schematic representation of A, illustrating the organization of hemangiosarcoma, osteosarcoma, and lymphoma xenografts. (**C**) Individual points on graph represent relative quantity of donor (dog) and host (mouse) cells in each tumor type. 10-12 fields of pictures at high magnification (400X) per slide were acquired. A total of approximately 1,000 cells in individual xenograft tumor was counted, and the percentages for each species-cells are presented.

To gain a deeper understanding of the involvement of stromal elements in canine hemangiosarcoma xenografts, we analyzed RNA-seq data obtained from cells and xenografts. The RNA-seq data was mapped to a reference genome that included both canine and mouse cells (26). Notably, we observed that the RNA-seq data from the cell lines predominantly mapped to the canine genome, whereas the xenografts exhibited mapping to both the canine and mouse genomes, which was consistent with the results obtained from FISH (**Supplementary Figure S2; Supplementary Table S2**).

### Canine xenografts generate lymphomas of mouse origin

An unexpected series of findings shed light on the relationship between hemangiosarcoma and inflammation in the microenvironment. Previous studies, including our own, have demonstrated that canine hemangiosarcoma cells can successfully form hemangiosarcomas in NSG mice (21–25). However, we noted instances where NSG mice, following inoculation with SB hemangiosarcoma cells or another cell line named Emma-brain (EFB), and BNX mice, following inoculation with tumor fragments derived from sample DHSA-1426 (freshly obtained from a dog with spontaneous hemangiosarcoma), developed round cell tumors. Notably, four out of five mice that received 5 × 10^6^ SB hemangiosarcoma cells intraperitoneally and one out of four mice that received 2 × 10^6^ EFB cells subcutaneously, succumbed acutely two weeks after inoculation, displaying signs of anemia and splenomegaly. Histological examination revealed that the spleens of these mice were expanded by monomorphic populations of hematopoietic cells (**Figure 3A, B**). Upon further analysis, the cells were determined to be of mouse origin and represented erythroid progenitors (Ter-119+), with a few canine hemangiosarcoma cells admixed in the population (**Figure 3C-F**). However, we were unable to definitively establish whether these cells had undergone malignant transformation.

**Figure 3.**
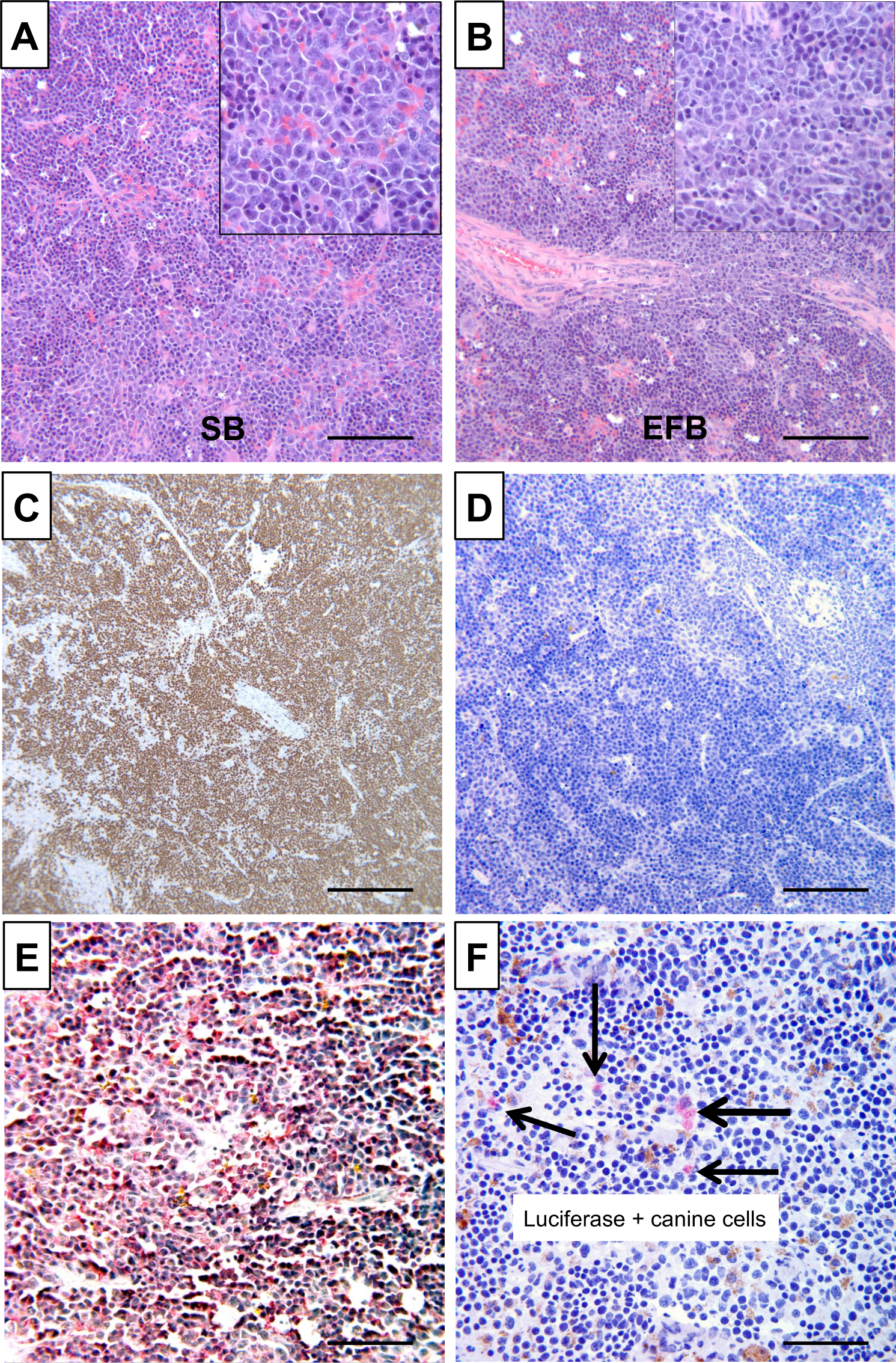
Hematopoietic expansion derived from adoptive transplantation of canine hemangiosarcoma in immunodeficient mice. (**A** and **B**) Xenotransplantation of canine hemangiosarcoma cell lines created exuberant myeloid hyperplasia in mouse spleens. Representative photomicrographs show histopathology by H&E staining of spleens from NSG mice transplanted with hemangiosarcoma cell lines SB (**A**) and EFB (**B**). (**C**) Immunoreactivity of anti-mouse Ki-67 (Tec-3) antibody shows strongly positive signal in proliferating cells in the spleen. (**D**) Immunostaining of anti-human (and canine cross-reactive) Ki-67 (MiB-1) antibody shows lack of positive staining among proliferating cells. (**E**) The proliferating cells are immunoreactive with anti-mouse Ter-119 antibody. (**F**) SB cells expressing Luciferase are detected in mouse spleen (arrows). Images shown in panels **C**, **D**, **E**, and **F** are from immunohistochemistry staining done in mice inoculated with SB cells modified to express GFP and firefly Luciferase. Horseradish peroxidase (for Ki-67 stains) and alkaline phosphatase (for Ter-119 and Luciferase) conjugates were used. Counterstain = hematoxylin. A-D: Bar = 200 µm; E-F: Bar = 50 µm.

Three of four mice that received DHSA-1426 tumor fragments, representing 1st-generation canine patient-derived xenograft (CPDX), developed tumors in multiple organs, including spleen, lymph nodes, meninges, cerebrum, and mesentery, 12 weeks after implantation. However, the morphology of these tumors resembled that of round cell tumors (**Supplementary Figure S3A, B**). Immunohistochemical analysis showed that the tumor cells expressed CD45, B220, and Pax5, but did not express CD3, Ter-119, and MPO (**Supplementary Figure S3C-H**), indicating a mouse B-cell origin. To further verify the mouse origin of these cells, flow cytometry was conducted, revealing positive staining for mouse CD45, but not for canine CD45 or human αVβ3-integrin (CD51/CD61), which exhibits cross-reactivity with canine, but not with mouse αVβ3-integrin (**Supplementary Figure S4**). Furthermore, RNA-seq data from the mouse round cell tumors demonstrated a near-complete alignment with the mouse genome, consistent with the findings from flow cytometry (**Supplementary Figure S2**), thereby indicating that the tumors were derived from mouse cells.

The malignant B-cells demonstrated the ability to undergo serial passage and establish B-cell lymphomas in recipient BNX mice independently, without the requirement for canine hemangiosarcoma cell support (**Supplementary Figure S5**). Intriguingly, similar results were obtained when single cell suspensions derived from fresh tumor fragments of canine hemangiosarcoma xenografts were inoculated into BNX recipients, representing 2nd generation CPDX derived from a 1st generation cell line tumor. The resulting tumors of mouse B-cell origin displayed comparable morphology and were likewise amenable to serial passage in BNX mice.

The potential etiology of these tumors was perplexing. To investigate the possibility of a transmissible, infectious agent as the cause of these tumors, we sequenced RNA from the xenografts and used the PathSeq platform for analysis. However, no bacterial or viral sequences with tumorigenic potential were identified in the xenografts or in the primary or metastatic canine hemangiosarcoma tumor samples or cell lines. While a recent study reported an association between *Bartonella spp*. and canine hemangiosarcoma (27,28), only 10 of 24 dogs tested in our study had detectable *Bartonella spp*. sequences, and these were present in low abundance (**Supplementary Table S3**). Furthermore, four of the samples contained sequences from *B. bacilliformis, B. grahamii*, or *B. tribocorum*, which infect humans and rats, respectively, and are not known to infect dogs as primary or accidental hosts (29,30). Three of the remaining six dogs had sequences for *B. clarridgeiae*, and three had sequences for multiple *Bartonella spp.*, including human and rodent-specific types. The low abundance and the presence of sequences from organisms that do not infect dogs suggest that the *Bartonella spp.* sequences were contaminants.

In contrast, murine leukemia virus (MuLV) reads were consistently identified in mouse B-cell lymphomas that arose from the hemangiosarcoma xenografts (**Supplementary Table S4**). The presence of MuLV sequences was further confirmed in one mouse B-cell lymphoma arising from hemangiosarcoma xenografts and in two subcutaneous canine hemangiosarcoma xenograft tumors through independent viral RNA sequencing experiments. Additionally, MuLV sequences were detected by polymerase chain reaction (PCR) in normal mouse tissues (liver and spleen), as well as in each of the three mouse B-cell lymphomas arising from the hemangiosarcoma xenografts and two subcutaneous canine hemangiosarcoma xenograft tumors. Notably, MuLV was also found in SB hemangiosarcoma cells that had been previously passaged through mice as hemangiosarcomas, suggesting that MuLV is a promiscuous virus with potential involvement in these tumors. These findings also suggested that MuLV might be the transforming agent responsible for the development of mouse lymphomas, and raised the possibility that canine hemangiosarcoma cells could provide a conducive environment for the expansion of these MuLV-transformed B-cells.

### Human and canine hemangiosarcoma tissues show evidence of immune cells

We reanalyzed RNA-Seq data obtained from both canine hemangiosarcoma and human angiosarcoma samples, with the aim of investigating the presence of immune cells within the tumor microenvironment. To identify co-expressed genes without bias, we used an unbiased approach called GCESS (Gene Cluster Expression Summary Score) (31). Notably, the highest expression of these immune signature genes was observed in the inflammatory subtype of canine hemangiosarcoma (**Supplementary Figure S6**). We next segregated human angiosarcomas into tumors with high and low immune scores (“immune-high” vs “immune-low”) and identified 461 up-regulated genes (FDR P < 0.05) in the immune-high group compared to immune-low group (**Figure 4A)**. Fifty-eight of these genes were also found in canine inflammatory hemangiosarcomas, and they were associated with T- and B-cell activation (**Figure 4B, C**). This approach allowed us to identify a common immune signature in both human and canine tumors.

**Figure 4.**
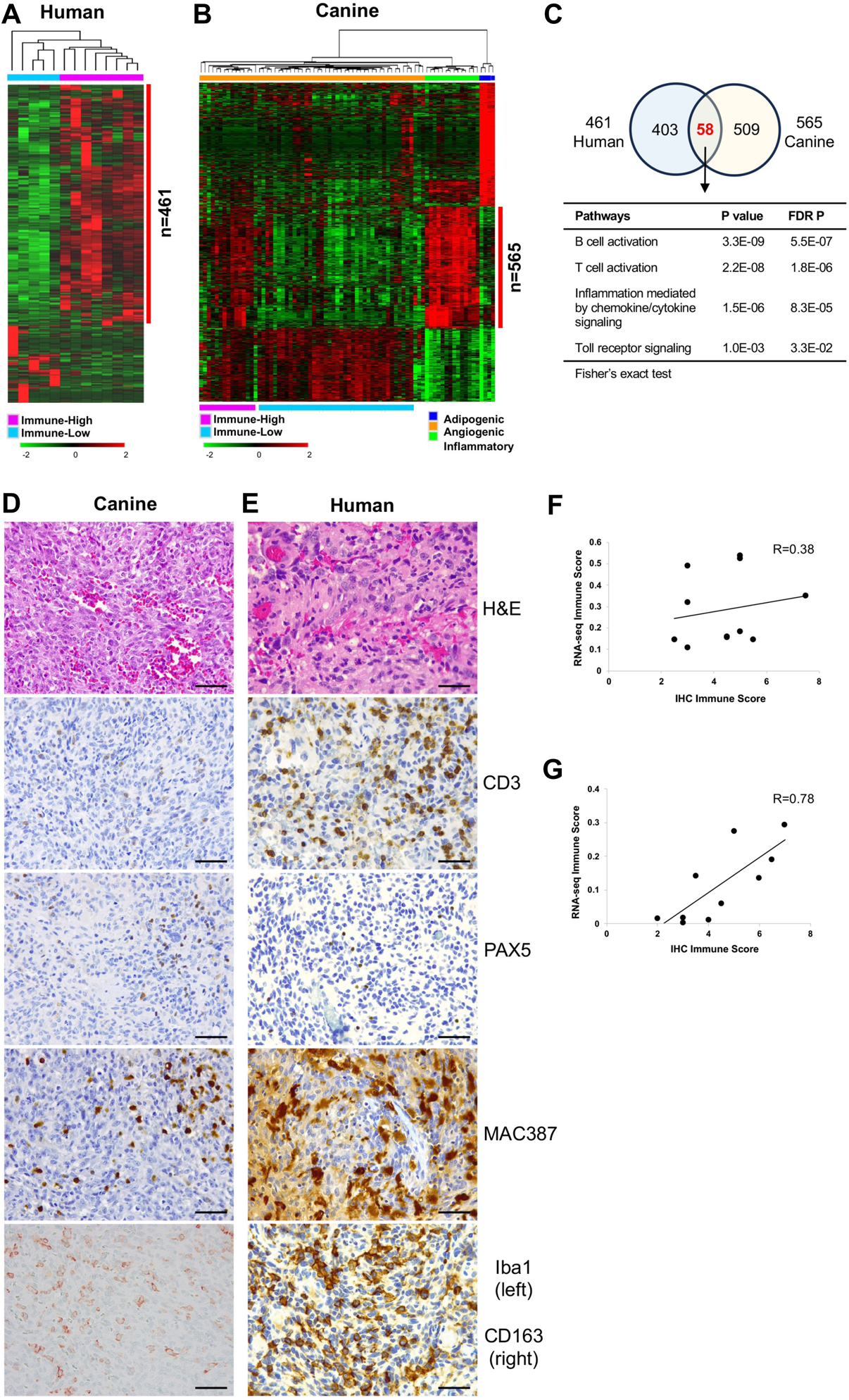
Immune cell infiltration and comparative immune signatures between canine hemangiosarcoma and human angiosarcoma. (**A**) 461 upregulated genes were identified in immune-high (N = 8) compared to immune-low (N = 5) groups in human angiosarcomas (FDR P value < 0.05). (**B**) 567 immune gene signatures were identified among three molecular subtypes of canine hemangiosarcomas (N = 76; FDR P value < 0.001; fold change > 3). The heatmaps show up-regulated (red) and down-regulated (green) genes by unsupervised hierarchical clustering (average linkage; mean-centered; log^2^ transformed). (**C**) Venn diagram shows 58 common genes associated with signaling pathways of immune cell functions between human and canine tumors. **(D** and **E)** Representative photomicrographs of H&E and immunohistochemical staining showing histological morphology and immune cell infiltration in canine hemangiosarcoma (**D**) and human angiosarcoma tissues (**E**) using anti-CD3, anti-PAX5, anti-MAC387, and anti-Iba1 (for canine) or anti-CD163 (for human) antibodies for detecting T cell, B cell, and macrophages. Horseradish peroxidase (counterstain = hematoxylin) or alkaline phosphate (for Iba1; counterstain = methylene blue). Bar = 50 um. (**F** and **G**) Scatter plots display correlation between transcriptional and immunohistochemical (IHC) immune score in canine hemangiosarcoma (**F**) and human angiosarcoma (**G**). Spearman’s correlation coefficient (R) was calculated.

To confirm whether the inflammatory gene signatures were originating from the malignant cells themselves or from inflammatory cells within the tumors, we employed bioinformatics tools and immunostaining techniques. Tumor purity was assessed using ESTIMATE and immune scores were assigned to each tumor using *xCell*. Both tools yielded consistent scores (**Supplementary Figure S7A, B**). As expected, we found a correlation between immune scores and the predominant transcriptional phenotype of the tumor in both canine and human samples (**Supplementary Figure S7C-F**). Specifically, angiogenic tumors exhibited low immune scores, whereas inflammatory tumors exhibited high immune scores.

To further validate that the observed immune signatures were the result of immune and inflammatory cells present within the tumors, we performed immunohistochemical staining on sections from 11 canine hemangiosarcomas and 10 human angiosarcomas. Antibodies against T cells (CD3), B cells (Pax5), myeloid cells (Mac387), and macrophages (Iba1 for canine; CD163 for human) were used; CD3+ T cells, Pax5+ B cells, Mac387+ myeloid cells, and Iba1+ or CD163+ macrophages were present in both canine and human tumors (**Figure 4D, E**). Myeloid cells were the most abundant, while T cells varied in frequency from rare to abundant, and B cells were infrequent. Importantly, the distribution of inflammatory cells was diffuse throughout the tumor tissue. Furthermore, there was a direct correlation between xCell immune scores and immunohistochemistry scores for both canine (Spearman R=0.38; P = 0.255) and human (Spearman R=0.78; P = 0.011) samples that were examined (**Figure 4F, G**). These findings provide evidence that the observed immune signatures in both canine and human tumors are likely attributed to the presence of immune and inflammatory cells within the tumor microenvironment.

### Canine hemangiosarcoma cells promote and maintain hematopoiesis

To determine whether hemangiosarcoma cells were functionally sufficient to expand hematopoietic progenitor cells *in vitro*, we conducted long-term culture initiating cell (LTC-IC) assays using DHSA-1426 and EFB canine hemangiosarcoma cells, to assess their potential in promoting and maintaining hematopoiesis in CD34+ human umbilical cord blood hematopoietic progenitor cells (HPCs). Mouse M2-10B4 and human bone marrow-derived mesenchymal stromal cells (MSCs) were used as positive controls, while HPCs cultured without feeder cells served as a negative control. Remarkably, DHSA-1426 cells were found to promote expansion of human CD34+ HPCs with at least comparable, if not superior, efficiency compared to conventional mouse or human feeder cells, and resulted in comparable proportions of hematopoietic cell differentiation *in vitro* across all lineages (**Figure 5; Supplementary Figure S8**). Similar results were observed with EFB cells, albeit with slightly more limited expansion and differentiation.

**Figure 5.**
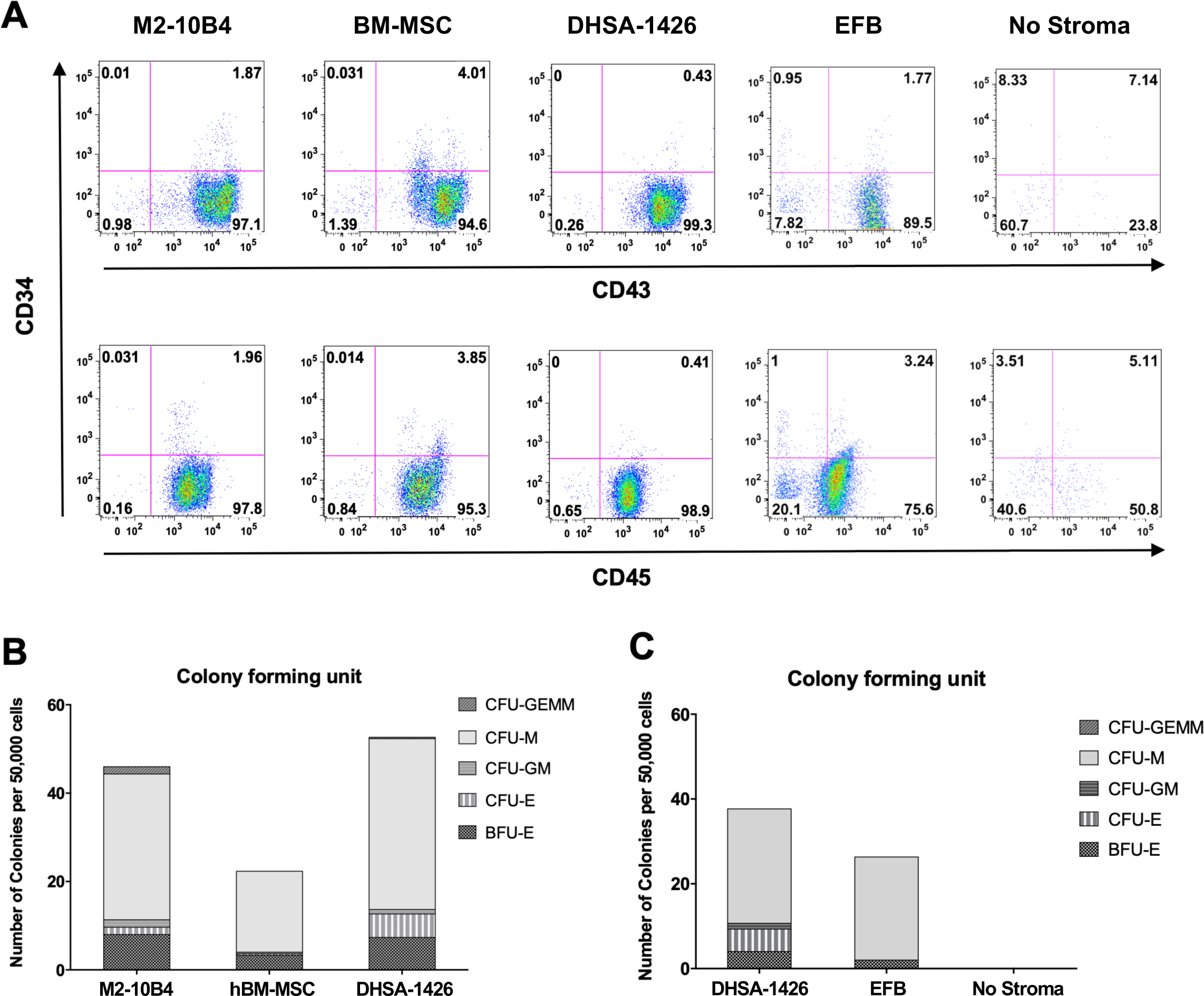
Canine hemangiosarcoma cells support expansion and differentiation of CD34+ human umbilical cord blood (hUCB) cells. (**A**) Flow cytometric data show populations of cells expressing CD43 and CD45 differentiated from CD34+ hUCB cells. CD34+ hUCB cells were pooled from two patients. M2-10B4, hBM-MSCs, and canine hemangiosarcoma cells (DHSA-1426 and EFB) were seeded on gelatin-coated 24-well plates at a density of 1×10^5^ cells/well. Gelatin-coated wells without stroma served as a negative control. Surface antigens of CD34, CD43, and CD45 were analyzed at week 5. (**B** and **C**) Bar graphs show number of different colonies formed by hUCB CD34+ cells co-cultured with feeder cells. Both DHSA-1426 and EFB canine hemangiosarcoma cell lines expanded hUCB CD34+ cells similar to the M2-10B4 and hBM-MSC positive control lines, while gelatin-coated wells alone failed to support expansion. Burst-forming unit-erythroid (BFU-E), CFU (colony-forming unit)-Erythroid (CFU-E), CFU-Granulocyte/Macrophage (CFU-GM), CFU-Macrophage (CFU-M), and CFU-Granulocyte/Erythroid/Macrophage/Megakaryocyte (CFU-GEMM) were determined for CFU assay.

## Discussion

Vasoformative sarcomas are aggressive tumors with uncertain cellular origin, proposed to arise from a multipotent bone marrow cell or lineage-committed endothelial progenitor cells in humans, dogs, and mice (1–4,8). While human angiosarcomas and canine hemangiosarcomas have a limited shared mutational spectrum, primarily in visceral forms of the disease and human breast angiosarcomas, convergent transcriptional programs characterized by deregulation of phosphoinositide 3-kinase pathways are activated in tumors of both species, as well as in zebrafish (8,32–38).

The transcriptional landscape of human angiosarcoma and canine hemangiosarcoma is strongly pro-angiogenic. However, a subset of tumors in both species exhibit robust transcriptional immune and inflammatory signatures that correlate with the presence of T cells and macrophages. The presence of immune and inflammatory infiltrates might be associated with longer survival outcomes, but further research is needed to determine their clinical significance regarding disease progression and their association with the approximately 15% of exceptional survivors reported in canine hemangiosarcoma (39,40).

The findings from our xenograft experiments provide further evidence supporting the concept that the disordered vascular organization in these tumors is driven by the tumor cells (1,21,25). Our experiments reveal that the tumor vessels are comprised of both malignant tumor cells and non-malignant host endothelial cells, a phenomenon that appears to be unique to hemangiosarcoma among the three types of xenografts we investigated. This observation underscores the ability of hemangiosarcoma cells to adopt endothelial functions and suggests that non-malignant cells play a role in the formation of aberrant blood vessels in vasculogenic tumors.

Interestingly, our experiments also demonstrate that the stromal cells in these xenografts contribute to the angiogenic and inflammatory transcriptional signatures. Our preliminary data further suggest that genes expressed by CAR cells, sinusoidal stromal cells, endosteal niche cells, endothelial progenitors, and hematopoietic progenitors (such as CSF3, IL6, IL11, and LIF) are enriched in canine hemangiosarcoma tissues and cells. This highlights the possibility that the stromal cells themselves are also influenced or reprogrammed by the malignant cells and supports a complex interplay between the tumor and its microenvironment. These findings highlight a potential vulnerability in the formation of the hemangiosarcoma niche, where the interactions between the tumor cells and their microenvironment are tightly orchestrated.

The initial perplexity surrounding the development of exuberant myeloid and erythroid hyperplasia, as well as bona fide lymphomas, arising from mouse cells in animals with primary or secondary hemangiosarcoma xenografts has led us to investigate the underlying etiology. Our results suggest that a transmissible etiology from dog to mouse is unlikely to be the cause of these expanded hematopoietic cell populations. Instead, our data indicate that canine hemangiosarcomas have the capability to support robust expansion and differentiation of hematopoietic progenitor cells *in vitro*, which may account for the expansion of myeloid cells *in vivo* in mice and in primary canine and human tumors. This expansion of non-malignant hematopoietic progenitor cells and mature leukocytes may also explain why clonal mutations (34) and fusions (8) are found less often in the inflammatory subtype of canine hemangiosarcoma.

In the case of lymphomas that occurred repeatedly and independently in animals receiving different tumor preparations, MuLV might have driven the transformation of residual lymphoid elements within a hyperproliferative environment created by the hemangiosarcoma cells. These findings are consistent with a previous report of angiosarcoma in the bone marrow of a human patient with tumor-associated myeloid proliferation and extramedullary hematopoiesis (41).

Other unexpected tumors have been reported in xenograft experiments and preclinical models of stem cell transplantation. For example, transplantation of murine MSCs has been shown to induce tumor formation and tissue malformation, possibly due to genetic instability and/or cellular transformation (42–44). Similarly, patient-derived xenografts of human solid cancers, such as breast, colon, pancreatic cancer, and rhabdomyosarcoma, have been reported to induce lymphomagenesis or lymphocytic tumors in immunodeficient mice, but in these cases, the tumors were derived from human tumor-infiltrating lymphocytes transformed by Epstein-Barr virus (45–48). Importantly, these previously reported tumors were all of donor origin, while the tumors observed in our study originated from the mouse recipients and were distinct from the donor hemangiosarcomas.

We were unable to find any reports of hematopoietic tumors of recipient origin arising from xenotransplantation experiments using other types of canine cancers in the literature, and we have not observed such events in our own studies (26,49,50). Thus, this finding appears to be unique to canine hemangiosarcoma and may be attributed to the ability of hemangiosarcoma cells to support hematopoietic expansion. Our findings also suggest that the normal counterparts of canine hemangiosarcoma cells might contribute to the development of hematopoietic malignancies through the creation of a permissive niche.

In this context, it is noteworthy that a shared region of the canine genome has been found to be significantly associated with B-cell lymphomas, hemangiosarcomas, and other blood derived tumors in various breeds of dogs belonging to distinct genetic clades (51–53). This finding highlights the intriguing possibility of a molecular connection between these seemingly distinct tumors, suggesting that disruptions in hematopoiesis and cellular reprogramming may contribute to their development (54). It also sets the origin of this potential genomic vulnerability prior to the derivation of modern dog breeds.

The capacity of the hematopoietic niche to accommodate bone marrow transplants and adoptive cell therapies has revealed its resilience, with bone marrow stromal cells exhibiting high resistance to chemotherapy and radiation. These intrinsic properties may explain the relatively poor long-term responses of human patients with angiosarcoma and dogs with hemangiosarcoma to cytotoxic therapies, and they could provide opportunities for the development of more effective treatments. However, it is important to recognize that therapies targeting the hematopoietic niche may also carry the potential for high toxicity.

Our data present a novel model to elucidate the etiology and cell of origin of canine hemangiosarcomas, and potentially human angiosarcomas. We propose that the malignant cells originate from bone marrow nurse cells, which have the capability to create niches that promote angiogenic proliferation or hematopoietic expansion, as illustrated in **Figure 6**. The robust inflammation observed in some of these tumors may therefore be intrinsic to the tumor itself, rather than solely due to extrinsic factors associated with tissue disruption. These paths of differentiation may also influence the biological behavior of the tumors, with those exhibiting strong angiogenic propensity displaying more aggressive behaviors (8).

**Figure 6.**
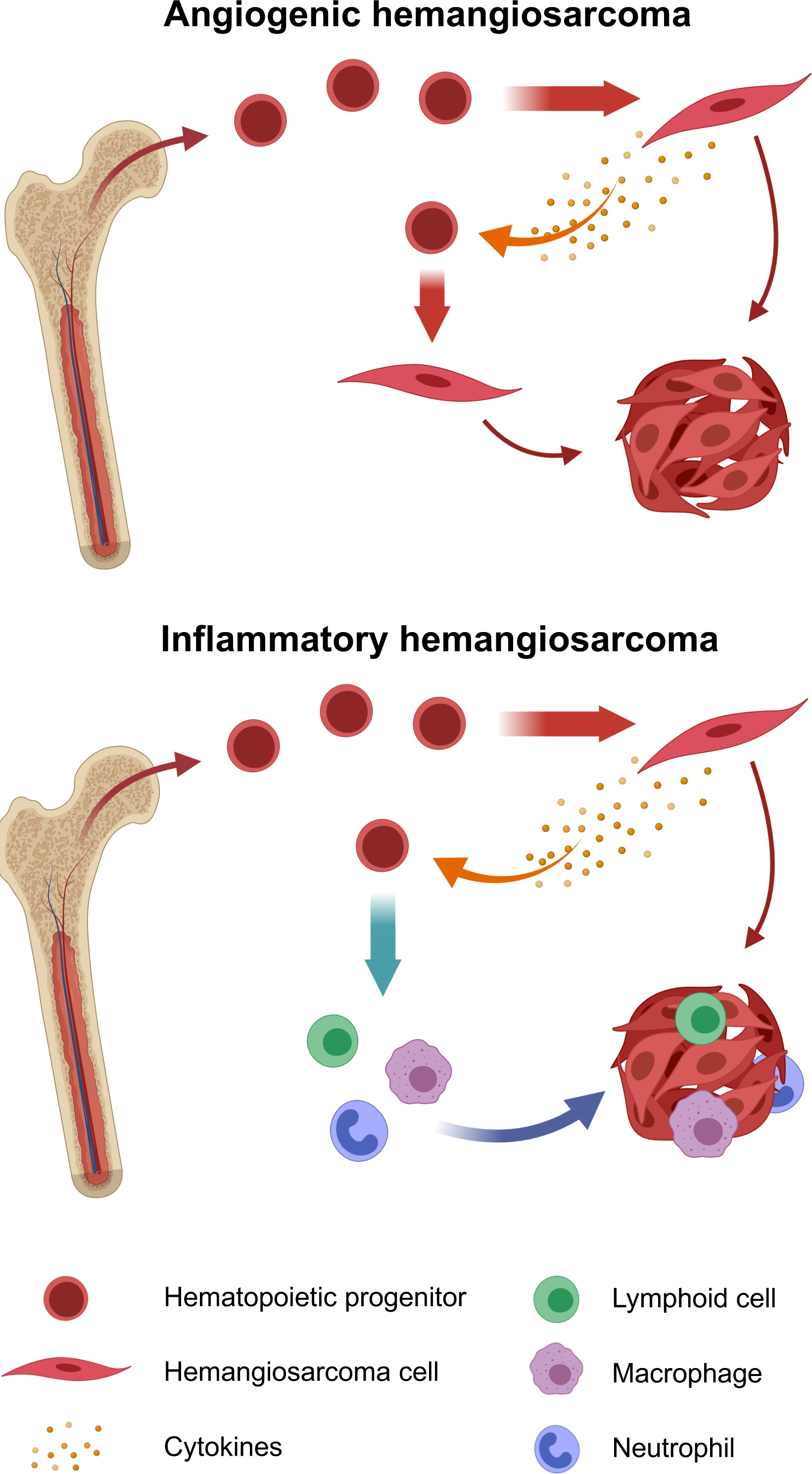
Hypothetical models for establishment of distinct molecular phenotypes of canine hemangiosarcoma. Non-mutually exclusive models illustrate discreet cells of origin for distinct molecular subtypes of hemangiosarcoma. Hemangiosarcoma may progress from bone marrow nurse cells that create a niche for hematopoietic expansion and inflammation, to a transitional pro-angiogenic state and full progression to a pure angiogenic state, or to malignant transformation with an inflammatory phenotype.

In conclusion, our study adds to the growing body of evidence demonstrating the complex interactions between tumor cells and the host microenvironment. Importantly, our data do not support a transmissible etiology for hemangiosarcoma. However, they do suggest that the permissive niche created by these cells can lead to the development of hematopoietic tumors driven by leukemia viruses in mice, raising the possibility that the bone marrow niche plays a similar role in viral lymphomas and leukemias in humans. Further investigations are warranted to unravel the underlying mechanisms that drive these observations and ascertain their significance in the development of human cancer.

## Materials and Methods

### Human tissue samples

The human angiosarcoma tissue samples used in this study were formalin-fixed and paraffin-embedded, as previously described (8). The samples were obtained from two sources: the University of Minnesota Biological Materials Procurement Network and the Cooperative Human Tissue Network. All sample acquisitions were performed under standardized patient consent protocols.

### Canine tissue samples and cell lines

Previously established hemangiosarcoma cell lines (SB, COSB, Emma, DD1, JHE, and JLU) (1,2,4,21,25,55) were used in this study. Canine tissue samples were collected from surgical removals or tumor biopsies at the University of Minnesota or private veterinary clinics. Additional hemangiosarcoma cell lines (DHSA-1401, DHSA-1420, and DHSA-1426) were generated and cultured using established methods (1,2,55). Cell line authentication for canine cells were conducted by IDEXX BioAnalytics (Columbia, MO, USA). All sample acquisition procedures were approved by the Institutional Animal Care and Use Committee (IACUC) of the University of Minnesota under protocols 0802A27363, 1101A94713, and 1312-31131A.

### Human and mouse cell lines

Human bone marrow-derived mesenchymal stromal cells (hBM-MSCs) were isolated from whole bone marrow purchased from AllCells (Emeryville, CA, USA), as described previously (56–59). M2-10B4 murine bone marrow stromal cells were purchased from the ATCC (Manassas, VA, USA) and maintained following established protocols (57). Samples of human umbilical cord blood (hUCB) were obtained from the ClinImmune Stem Cell Laboratory at the University of Colorado Cord Blood Bank (60).

### Mice and xenotransplantation

We conducted multiple xenograft experiments using various cell lines and patient-derived tumor sections. First, we injected cultured-tumor cells from three hemangiosarcoma cell lines (SB, Emma, and JHE) into NSG mice. Specifically, we injected 5 × 10^6^ SB cells subcutaneously in four mice, and 5 or 10 × 10^6^ SB cells intraperitoneally in four mice. Additionally, we injected 2 × 10^6^ Emma cells subcutaneously and intraperitoneally in four mice each, and 3 × 10^5^ JHE cells subcutaneously in four mice. Next, we injected tumor cells from five hemangiosarcoma cell lines (Emma, DD1, JLU, DHSA-1401, and COSB) into NSG neonates, 1 or 2 days after birth. We injected 5 × 10^5^ Emma, DD1, JLU, or DHSA-1401 cells, or 6.25 × 10^5^ COSB cells intraperitoneally in 50 µL of PBS. We also injected 5 × 10^6^ cells from three hemangiosarcoma cell lines (JLU, DHSA-1420, and DHSA-1426) in a mixture of 100 µL of PBS and 100 µL of BD Matrigel™ Basement Membrane Matrix into the subcutaneous space of BNX mice. For DHSA-1426, we injected tumor cells from passage-5 and passage-14 independently. We used sections of viable tumors from four dogs affected with hemangiosarcoma and implanted them into subcutaneous pockets of four mice for each dog. In addition, we implanted sections of non-hemangiosarcoma splenic tissues from seven dogs into 18 mice as controls. Finally, after visible tumors developed in mice, we serially transplanted the tumors by inoculation of cultured tumor cells in three mice or by direct implantation of single-cell suspensions of the tumor in eight mice. We monitored the mice for tumor development and sacrificed them when they reached a tumor endpoint, including a mass measuring 1.5 cm in the longest diameter or at the end of a 16-week period after xenotransplantation. All animal procedures were conducted in accordance with the Research Animal Resources (RAR) husbandry and care protocols and reviewed and approved by the IACUC of the University of Minnesota (protocols 1006A84813, 1106A00649, 1306-30712A, and 1311-31104A). A total of 132 mice were used for xenotransplantation procedures, as described in **Supplementary Table S1**.

### Flow cytometric analysis

We used the following antibodies for flow cytometry analysis of xenograft tumors: rat anti-dog CD45-APC (Clone YKIX716.13; AbD Serotec, Raleigh, NC, USA), rat anti-mouse CD45-APC (Clone 30-F11; eBioscience, San Diego, CA, USA), and mouse anti-human αvβ3 (CD51/CD61)-FITC (Clone 23C6; BD Pharmingen, San Jose, CA, USA). Isotype controls included rat IgG2b-APC (Clone eB149/10H5; eBioscience) and mouse IgG1 K-FITC (Clone P3.6.2.8.1; eBioscience). Data were acquired using a BD Acuri C6 flow cytometer (BD Biosciences, San Jose, CA) and analyzed with FlowJo software (version 10.1, Tree Star, Ashland, OR, USA).

### Immunohistochemistry

Immunohistochemical staining was conducted at either the BioNet Histology Laboratory, Minnesota Veterinary Diagnostic Laboratory, or Comparative Pathology Shared Resource at the University of Minnesota. The following antibodies were used: CD3, PAX5, MAC387, CD163, CD204, Iba1, Ter-119, MPO, and CD45. Details of the antibodies used are provided in **Supplementary Table S5**.

### RNA-seq for gene expression profiling

To obtain transcriptomic profiles, we isolated total RNA from xenograft tumor samples using the TriPure Isolation Reagent (Roche Applied Science, Penzberg, Germany), followed by clean-up using the RNeasy Mini Kit (Qiagen). We generated RNA-seq libraries with a targeted depth of 20 million paired-end reads (2 x 50 bp) at the UMGC, as previously described (4,8). We conducted bioinformatic analyses for gene expression profiling as previously reported (4,8,26,31,61). RNA-seq data from human angiosarcoma tissues can be accessed via the Gene Expression Omnibus (GEO) under accession number GSE163359 (8). RNA-Seq data from canine hemangiosarcoma tissues are available through the GEO (accession number GSE95183) and the NCBI Sequence Read Archive (accession number PRJNA562916) (8).

### Viral RNA isolation and library generation

To detect RNA viruses, we isolated viral RNA from xenograft tumors using either a Qiagen viral RNA kit or Trizol followed by the Qiagen viral RNA kit (Qiagen, Germantown, MD, USA). The isolated RNA was then submitted to the University of Minnesota Genomics Center (UMGC) for viral RNA MiSeq. The Illumina MiSeq generated approximately 1 million paired-end reads (2 x 250) for each sample.

### PathSeq analysis

The PathSeq algorithm was used to determine transmissible agents in RNA-seq data from dog and mouse samples. The PathSeq algorithm (62) was used to perform computational subtraction of mouse and dog genomes followed by alignment of residual reads to mouse and dog reference genomes and microbial reference genomes (including bacterial, viral, archaeal, and fungal sequences downloaded from NCBI). These alignments resulted in the identification of microbial reads in the data. Human reads were subtracted by first mapping reads to a database of mouse (mm9) and dog reference genomes (canFam3.1; GCF_000002285) using BWA (Release 0.6.1, default settings) (63), MegaBLAST (Release 2.2.25, cut-off E-value 10^-7^, word size 16) and BLASTN (Release 2.2.25, cut-off E-value 10^-7^, word size 7, nucleotide match reward 1, nucleotide mismatch score -3, gap open cost 5, gap extension cost 2) (64). Only sequences with perfect or near perfect matches to the human genome were removed in the subtraction process. In addition, low complexity and highly repetitive reads were removed using RepeatMasker (version open-3.3.0) (65). To identify microbial reads, the residual reads were aligned with MegaBLAST to a database of microbial and dog reference genomes. Raw read counts were calculated using the reads that were mapped to *Epstein-Barr virus* (EBV) and *H. pylori* with at least 90% identity and 90% query coverage. Using the raw read counts, the abundance metric or normalized read count of a given microbe in a sample was calculated as

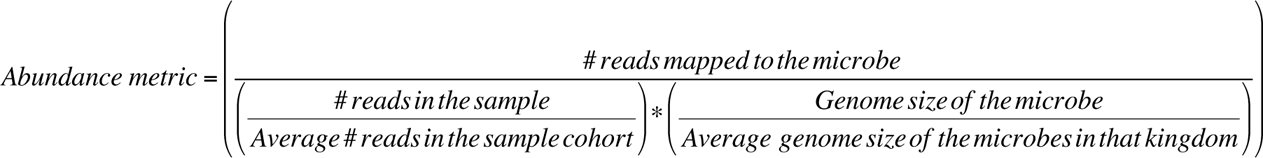

Relative abundance in a given sample was calculated as abundance metric of taxa divided by the total abundance metric at kingdom level of the sample.

### Long-term culture initiating cell (LTC-IC) and hematopoietic colony forming unit (CFU) assays

We isolated human umbilical cord blood (hUCB) CD34+ cells from two patients using the Miltenyi CD34 Microbead Kit and MACS separation column (Miltenyi Biotec, Boston, MA, USA), following the manufacturer’s instructions. The purity of isolated hUCB CD34+ cells was determined by flow cytometry using the following antibodies (all anti-human): CD34-PECy7 (eBioscience, San Diego, CA, USA), CD43-APC (BD Biosciences, San Jose, CA, USA), and CD45-APC (BD Biosciences). Sorted hUCB CD34+ cells with >92% CD34+CD45+ were used for subsequent experiments. We seeded M2-10B4, hBM-MSCs, 1426, and Emma in gelatin-coated 24-well plates at a density of 10^5^ cells/well overnight. After inactivation with 10 μg/mL Mitomycin C for three hours, 5,000 hUCB CD34+ cells were added to each well in Myelocult H5100 (StemCell Technologies, Vancouver, BC, Canada) supplemented with 1μM dexamethasone (Sigma-Aldrich, St. Louis, MO, USA). Gelatin-coated wells without stroma were used as negative controls. Cells were conditioned for six weeks at 37°C and 5% CO_2_ with one-half media exchanges every week. At week 5, non-adherent cells from selected wells were harvested, counted, and analyzed for hematopoietic-specific surface antigens using the following antibodies (all anti-human): CD34-PECy7, CD33-APC (BD Biosciences), CD43-APC and CD45-APC. At the end of the six-week culture, both non-adherent and adherent cells were collected using 0.05% trypsin supplemented with 2% chicken serum for five minutes. Cells were filtered using a 70 μm cell strainer to generate single-cell suspensions and counted. We prepared triplicate CFU assays by seeding 50,000 cells/1.5 mL of semi-solid Methocult H4435 Enriched media (StemCell Technologies) in 35 mm culture dishes (Greiner, Monroe, NC, USA). After two additional weeks, the plates were manually scored for total number and phenotype of the CFUs, as previously described (66).

### Statistical analysis

Pearson and Spearman correlation coefficient was calculated for correlation between two variables. Statistical analysis was performed using GraphPad Prism 9 (GraphPad Software, Inc., San Diego, CA, USA) or Microsoft Excel.

### Data Availability

The data generated in this study are publicly available in GEO at GSE163359 and GSE95183, as well as in NCBI Sequence Read Archive (PRJNA562916). Other relevant data generated in this study are available upon request from the corresponding author.

### Authors’ Contributions

Conception and design: J.H. Kim, J.F. Modiano

Development of methodology: J.H. Kim, A.L. Sarver, M.G. Angelos, A.M. Frantz, S. Bullman, C.S. Pedamallu, S. Kumar, S.M. Goyal, L. Oseth, C.L. Foster, T.D. O’Brien, I. Cornax, J.F. Modiano

Acquisition of data (provided animals, acquired and managed patients, provided facilities, etc): M.G. O’Sullivan, M. Meyerson, T.C. Lund, J. Alfoldi, M. Breen, K. Lindblad-Toh, D.S. Kaufman, J.F. Modiano

Analysis and interpretation of data (e.g., statistical analysis, biostatistics, computational analysis): J.H. Kim., A.L. Sarver., M.G. Angelos, A.M. Frantz, N. Cheng., J. Alfoldi., K. Lindblad-Toh, S. Bullman, C.S. Pedamallu

Writing, review, and/or revision of the manuscript: J.H. Kim, A.J. Schulte, E.B. Dickerson, D.S. Kaufman, J.F. Modiano

Administrative, technical, or material support (i.e., reporting or organizing data, constructing databases): J.H. Kim, A.J. Schulte, M. Lewellen.

Study supervision: J.H. Kim, J.F. Modiano

## Acknowledgements

The authors acknowledge Keumsoon Im for assistance with experiments and data acquisition. The authors would also like to acknowledge Milcah Scott for processing the next generation sequencing data and assistance with data analysis. Artistic design of Figures 2A and 7 was created with Biorender (biorender.com). This work was partially supported by grants 1R03CA191713-01 (to J.F. Modiano, A.L. Sarver, and J.H. Kim) from the NCI of the NIH, grants #02759 (to J.H. Kim), #422 (to J.F. Modiano), and 1889-G (to J.F. Modiano, M. Breen, and K. Lindblad-Toh) from the AKC Canine Health Foundation, grant JHK15MN-004 (to J.H. Kim, A.L. Sarver, J.F. Modiano) from the National Canine Cancer Foundation, grant D10-501 (to J.F. Modiano, M. Breen, and K. Lindblad-Toh) from Morris Animal Foundation, and grant from Swedish Cancerfonden (to K. Lindblad-Toh). This work was supported by an NIH NCI R50 grant, CA211249 (to A.L. Sarver). It was also supported by a Career Development Award from the Department of Defense Peer Reviewed Cancer Research Program (to J.H.Kim). The NIH Comprehensive Cancer Center Support Grant to the Masonic Cancer Center, University of Minnesota (P30 CA077598) provided support for the cytogenetic analyses performed in the Cytogenomics Shared Resource. M. Breen is supported in part by the Oscar J. Fletcher Distinguished Professorship in Comparative Oncology Genetics at North Carolina State University. K. Lindblad-Toh is supported by a Distinguished Professor award from the Swedish Research Council. J.F. Modiano is supported by the Alvin and June Perlman Chair in Animal Oncology. The authors gratefully acknowledge donations to the Animal Cancer Care and Research Program of the University of Minnesota that helped support this project. The content of this manuscript is solely the responsibility of the authors and does not necessarily represent the official views of any of the funding agencies listed above.

