## Supplementary Figures for "Hemangiosarcoma Cells Promote Conserved Host-Derived Hematopoietic Expansion": Supplementary_Figs.pdf

### Supplementary Figure S1

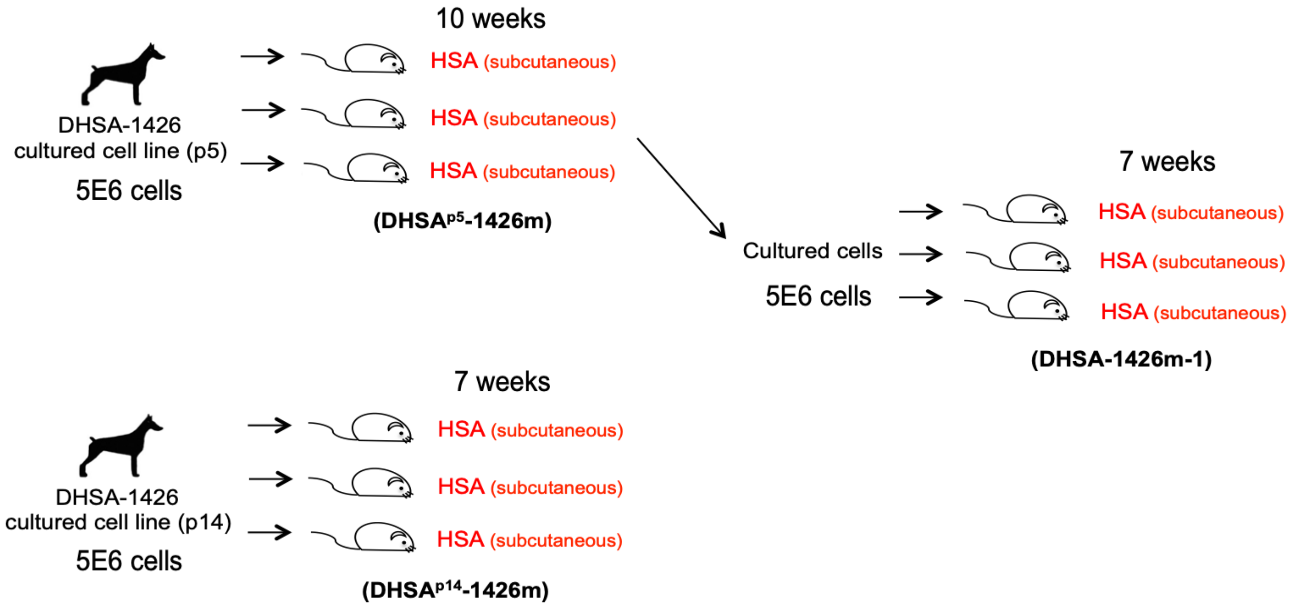


**Supplementary Figure S1. Serial transplantation of canine hemangiosarcoma cells in immunodeficient mice.** Xenograft tumors were reproducible by inoculation of passage 5 (p5) and p14, DHSA-1426 cells. DHSA-1426 cells from p5 were serially passaged into new recipient mice and tumor development was accelerated from 10 weeks in the initial xenografts to seven weeks in the first passage xenografts. The additional time in culture (from p5 to p14) also accelerated development of xenografts from 10 to seven weeks.

#### Supplementary Figure S2

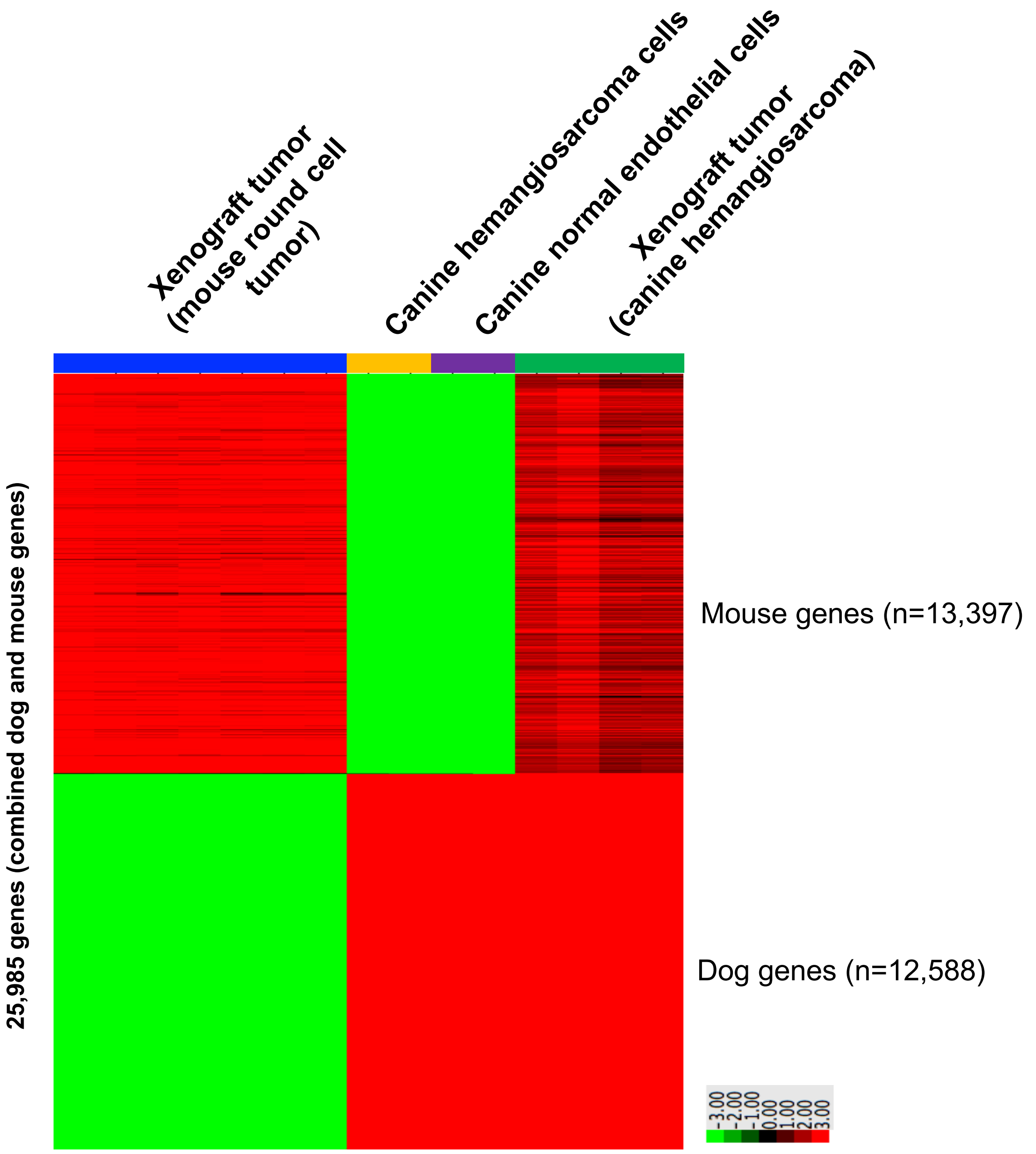


**Figure S2. Identification of dog tumor and mouse stromal signatures from hemangiosarcoma xenografts.** We identified donor (dog) and host (mouse) genes in mouse round cell tumors (N = 7; blue column bar) and hemangiosarcoma xenografts (N = 4; green column bar) using RNA-seq and our dog/mouse hybrid genome bioinformatics pipeline. Sequencing reads to canine and murine genes within xenograft tumors were mapped to dog (canFam3) and mouse reference genome (mm10) using a HISAT2. Species-specific gene counts were calculated as described. Canine hemangiosarcoma cells (N = 2; yellow column bar) and normal endothelial cells (CnAOEC; N = 2; purple column bar) cultured in *in vitro* were used as controls. A heat map shows up-regulated (red) and down-regulated (green) genes between genes by unsupervised hierarchical clustering (average linkage; mean- centered; log2 transformed).

### Supplementary Figure S3

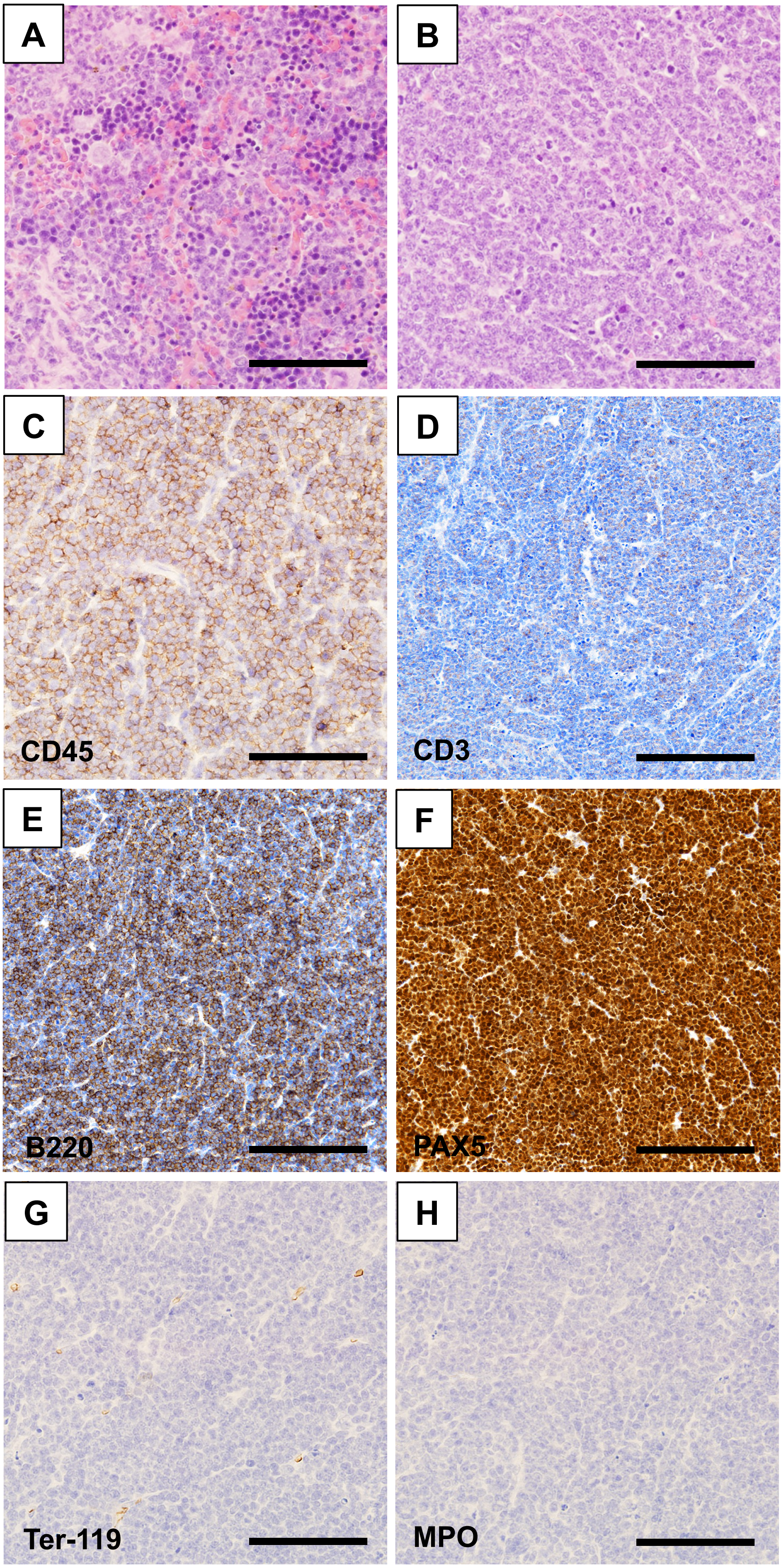


**Figure S3. Round cell tumor developed from xenotransplantation of canine hemangiosarcoma in immunodeficient mice.** (**A** and **B**) Representative photomicrographs display histopathology of tumors from BNX mice transplanted with DHSA-1426 hemangiosarcoma cell line. H&E stain; splenic tumor (**A**) and tumor in lymph node (**B**). Immunohistochemistry was done in lymph node tumors with anti-CD45 (**C**), anti-CD3 (**D**), anti-B220 (**E**), anti-PAX5 (**F**), anti-Ter-119 (**G**), and anti-MPO (**H**) antibodies. Horseradish peroxidase conjugates were used. Counterstain = hematoxylin. Bar = 50 µm.

### Supplementary Figure S4

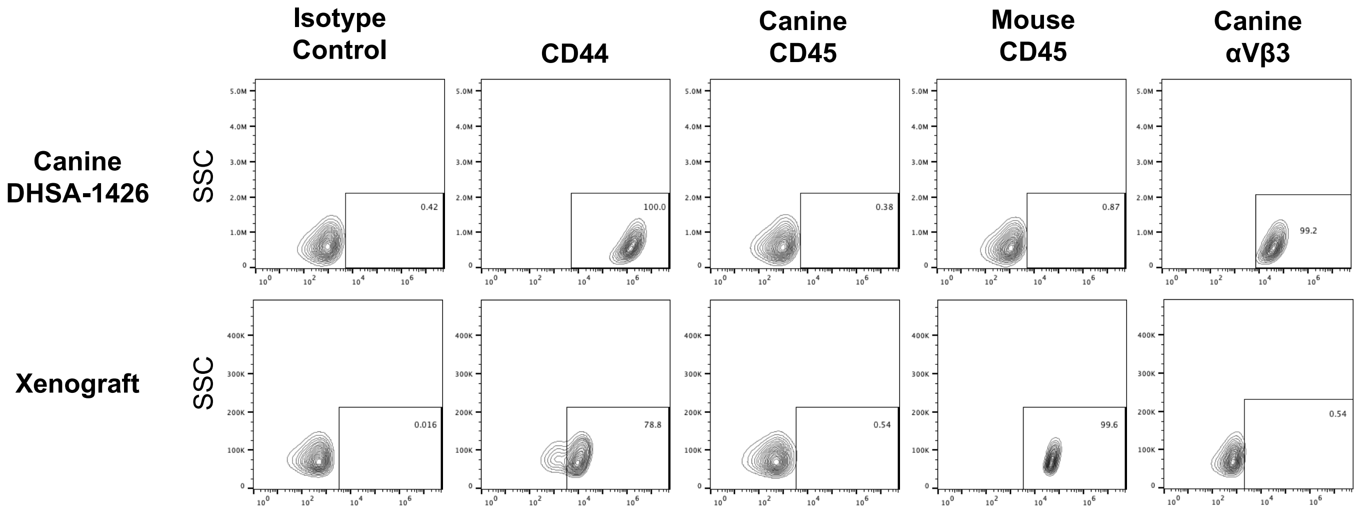


**Summary of staining**

| | CD44 | Canine CD45 | Mouse CD45 | Canine αVβ3 |
|---|---|---|---|---|
| **DHSA-1426 cells** | High | Low | Low | High |
| **Xenograft tumor cells** | High | Low | High | Low |

**Figure S4. Flow cytometric analysis of mouse round cell tumors from xenograft of canine hemangiosarcoma**. Contour plots show cell population stained with anti-canine CD45, anti-mouse CD45, and anti-human αVβ3-integrin antibodies in canine DHSA-1426 cells and mouse round cell tumors. Anti-CD44 antibody (dog- and mouse-cross-reactive) was used as staining positive control.

### Supplementary Figure S5

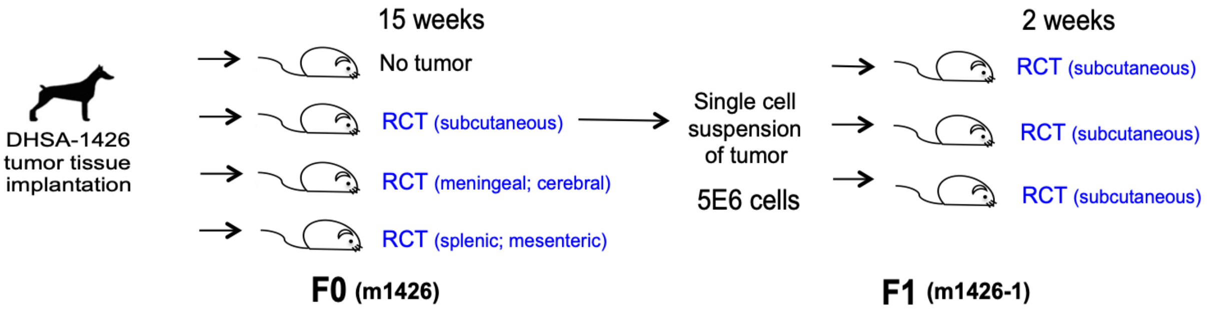


**Figure S5. Schematic illustration of incidence of mouse round cell tumor and hemangiosarcoma in xenotransplantation of canine hemangiosarcoma in immunodeficient mice.** Splenic tumor fragment obtained from a dog with hemangiosarcoma was surgically implanted into the flank of BNX mice. Three of four mice developed round cell tumors at 15 weeks. After the tumors were harvested, $5x10^6$ tumor cells in single cell suspension were implanted into three mice, and all of the mice developed round cell tumors two weeks later.

#### Supplementary Figure S6

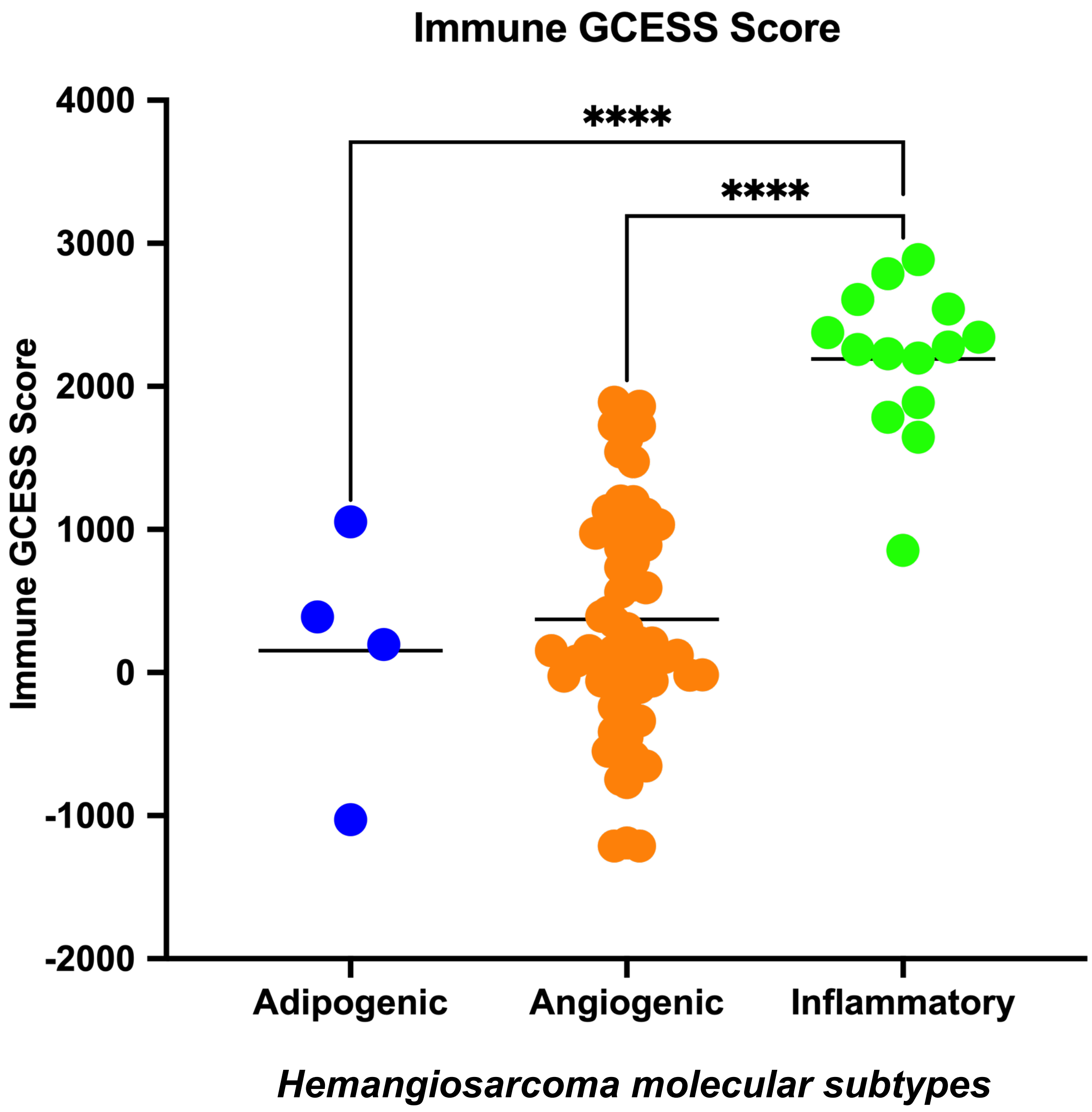


**Figure S6. Immune GCESS scores in RNA-seq data of canine hemangiosarcoma tissues.** (**A**) A total of 76 canine hemangiosarcomas were analyzed for immune GCESS scores (adipogenic, N=4; angiogenic, N=58; inflammatory, N=14). Three molecular subtypes of canine hemangiosarcomas were identified as previously described [Ref 4,8]. One-way ANOVA test; ****, P<.0.0001

### Supplementary Figure S7

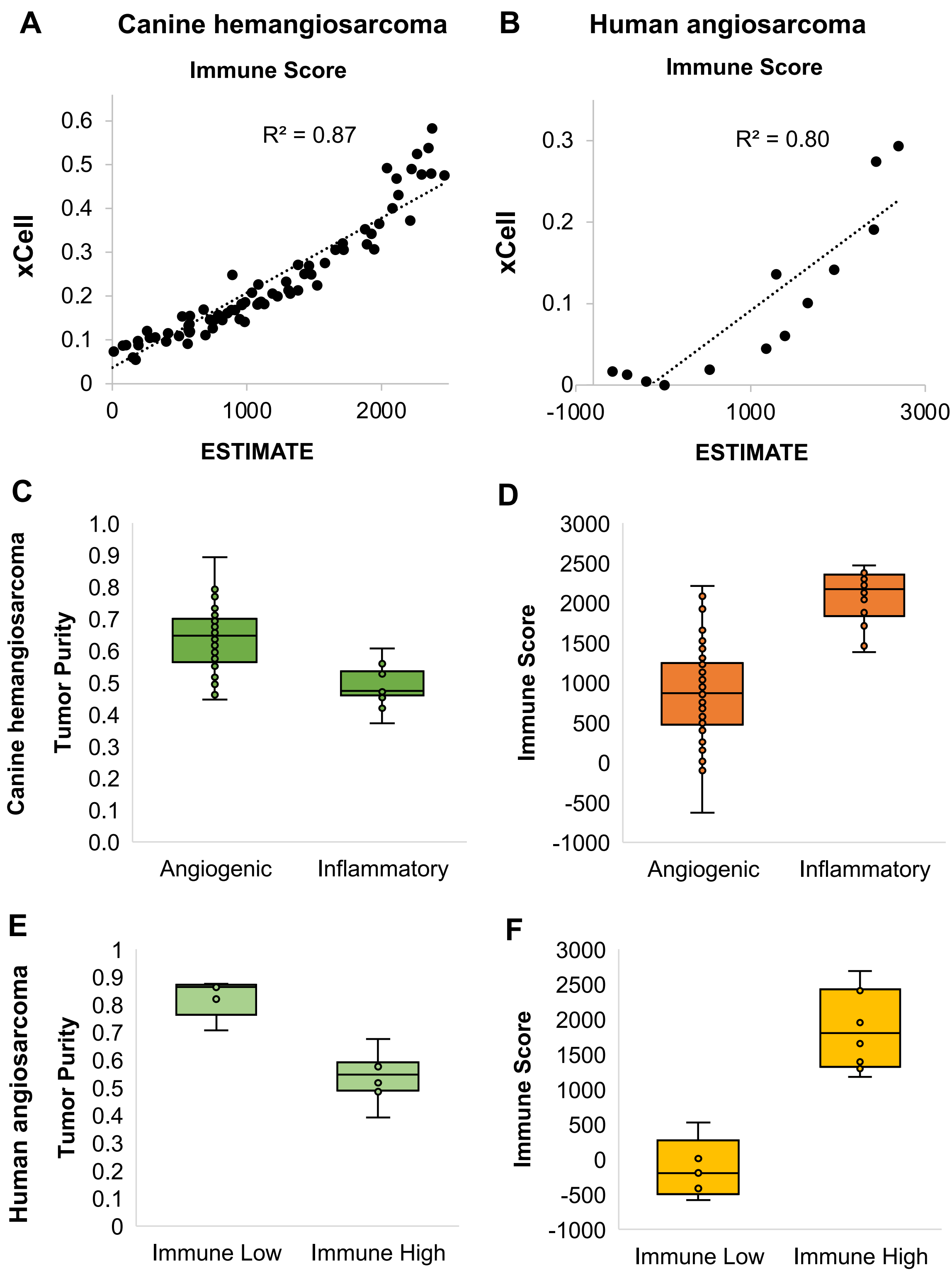


**Figure S7. Transcriptional immune and tumor purity score in canine hemangiosarcoma and human angiosarcoma.** (**A** and **B**) Scatter plots display correlation between immune scores predicted by ESTIMATE (x-axis) and *xCell* (y-axis) tools in canine hemangiosarcoma (**A**) and human angiosarcoma (**B**). Coefficient of determination (R2) was calculated by linear regression. (**C** - **F**) Box and Whisker plots show tumor purity and immune scores between canine angiogenic and inflammatory hemangiosarcoma (**C** and **D**) and between human angiosarcoma with low and high immune signature (**E** and **F**).

#### Supplementary Figure S8

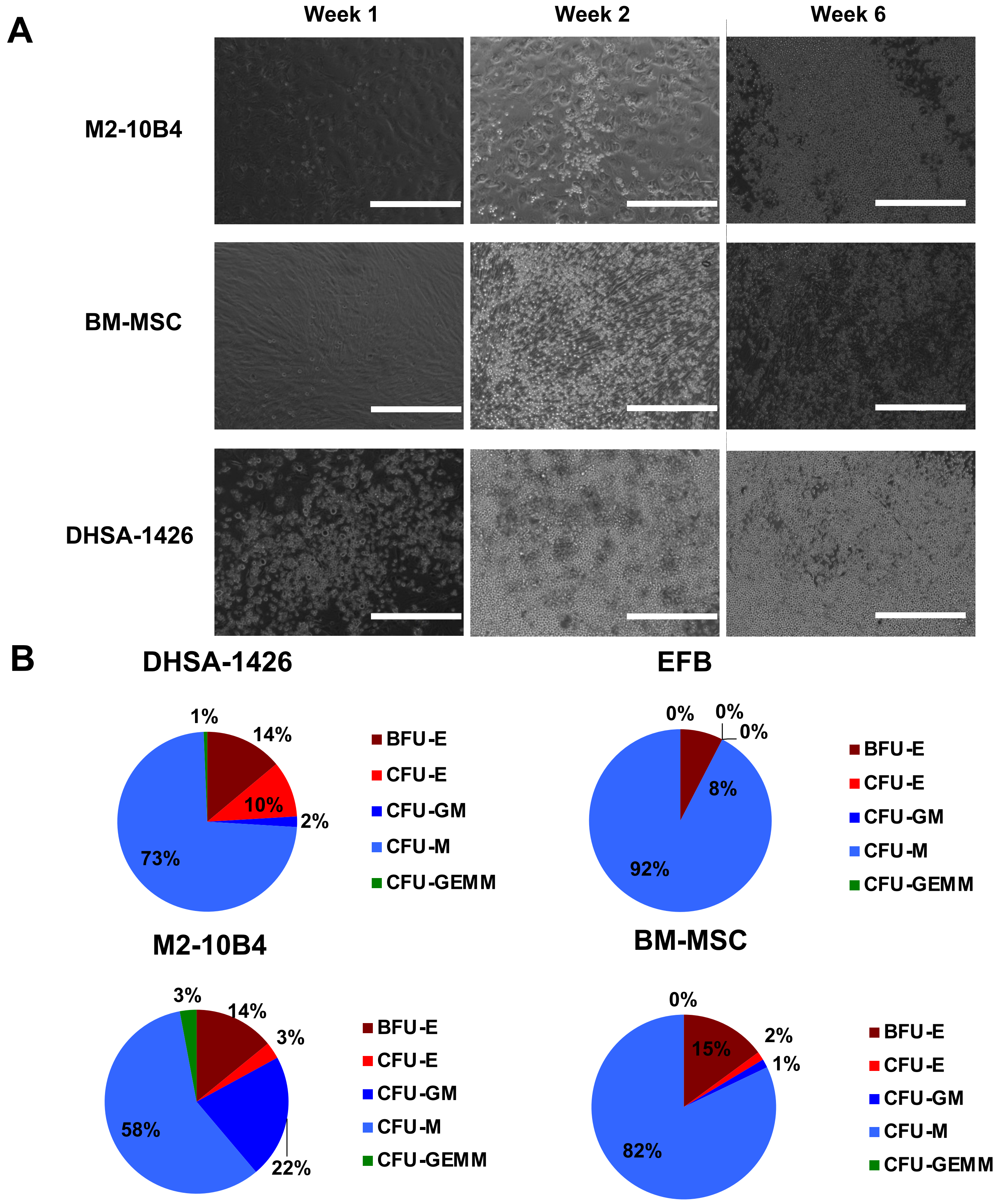


**Figure S8. Long-term culture initiating cell and hematopoietic colony-forming unit assays on CD34+ cells and canine hemangiosarcoma cells. (A)** Representative photomicrographs display proliferation of CD34+ hUCB cells co-cultured with M2-10B4, BM-MSC, and DHSA-1426 cells at week 1, 2 and 6. Bar = 400 µm. (**B**) Pie charts present colony-forming units differentiated from CD34+ hUCB cells by co-culture with different feeder cells, DHSA-1426, EFB, M2-10B4, and BM-MSC.
